# Investigating the relation between stochastic differentiation and homeostasis in intestinal crypts via multiscale modeling

**DOI:** 10.1101/000927

**Authors:** Alex Graudenzi, Giulio Caravagna, Giovanni De Matteis, Marco Antoniotti

## Abstract

Colorectal tumors originate and develop within intestinal crypts. Even though some of the essential phenomena that characterize crypt structure and dynamics have been effectively described in the past, the relation between the differentiation process and the overall crypt homeostasis is still partially understood. We here investigate this relation and other important biological phenomena by introducing a novel multiscale model that combines a morphological description of the crypt with a gene regulation model: the emergent dynamical behavior of the underlying gene regulatory network drives cell growth and differentiation processes, linking the two distinct spatio-temporal levels. The model relies on a few a priori assumptions, yet accounting for several key processes related to crypt functioning, such as: dynamic gene activation patterns, stochastic differentiation, signaling pathways ruling cell adhesion properties, cell displacement, cell growth, mitosis, apoptosis and the presence of biological noise.

We show that this modeling approach captures the major dynamical phenomena that characterize the regular physiology of crypts, such as cell sorting, coordinate migration, dynamic turnover, stem cell niche maintenance and clonal expansion. All in all, the model suggests that the process of stochastic differentiation might be sufficient to drive the crypt to homeostasis, under certain crypt configurations. Besides, our approach allows to make precise quantitative inferences that, when possible, were matched to the current biological knowledge and it permits to investigate the role of gene-level perturbations, with reference to cancer development. We also remark the theoretical framework is general and may applied to different tissues, organs or organisms.

## Introduction

Intestinal crypts are invaginations in the intestine connective tissue, which are the *loci* where *colorectal tumors*, one of the major causes of deaths in adults, originate and develop [1–4]. These particular structures have been quite precisely characterized, highlighting a fast renewing single layer epithelium in which distinct cell populations are rather sharply stratified and cells coordinately migrate from the *multi-potent stem cell* niche (at bottom) toward the intestinal lumen, with some exceptions [5–9]. As long as cells move upward they divide and differentiate through intermediated stages, according to a hypothesized *lineage commitment tree*, which ensures the correct functioning of the crypt and its resistance to perturbations and biological noise. The complex interplay between *cell proliferation, differentiation, migration* and *apoptosis* results in the overall homeostasis of the system. Chemical gradients ruled by key signaling pathways such as *Wnt, Notch, Eph/ephrin* have a crucial role in all these processes and, when progressively mutated or alterated, cancerous structures may emerge [10, 11].

Mathematical and computational models have been widely used to describe intestinal crypts (see [12, 13] and references therein). Among these, *compartmental models* analyze population dynamics via mean-field approaches without accounting for the spatial and mechanical properties of the crypts [14, 15]. In order to consider *space*, both *in-lattice* and *off-lattice* models have been defined. The former use simplified cellular automata-based representations of crypts to account for cell displacement, movement and interactions (see, e.g., [16, 17]). The latter strive to model more directly the geometry and the physics of crypts, but, as they involve bio-mechanical forces and complex geometries (e.g., *Voronoi diagrams*), the spaces of parameters and variables dramatically enlarges (see, e.g., [18–20]). As usual, the best trade-off between the complexity of the model and that of the modeled phenomena depends on the aim of the research.

Even if a large list of important phenomena, such as the spatial arrangement of cell population or the stem cell niche maintenance, have been described with noteworthy results with currently existing models, the relation between the underlying differentiation processes and the overall crypt homeostasis is still partially understood. To investigate *in-silico* this relation and other important biological properties we here introduce a *novel* multiscale model of intestinal crypt dynamics. The *multiscale* approach allows to consider, at different abstraction levels, phenomena happening at distinct spatiotemporal scales, as well as the hierarchy and the communication rules among them [21, 22]. In the case of crypts, these include intra-cellular processes such as gene regulation and intra-cellular communication, and inter-cellular processes such as signaling pathways, inter-cellular communication and microenvironment interactions. Their joint complex interaction allows to quantify, at the level of *tissues*, some key properties of crypts such as their spatial patterning, cellular movement, migration and homeostasis.

The foundations of our model lay in *statistical physics* and in *complex systems theory*, as the main rationale is to use the simplest possible model to reproduce relevant complex phenomena, also allowing for a comparison with experimental data and biological knowledge [23]. Thus, our model relies on few *a priori* assumptions and constraints, and most of its properties are *emergent*. The model is composed of two distinct levels, accounting for the crypt *morphology* and the underlying cellular *Gene Regulatory Network* (GRN).

Crypt morphology, the spatial level of the model, is described via the well-known *in-lattice Cellular Potts Model* (CPM), already proven to reproduce several properties of real systems [24–26]. In this discrete representation cells are represented as contiguous lattice sites (i.e. *pixels*), and their movement (via pixel re-assignment) is driven by an energy minimization criterion accounting for cellular type, position, age and size. Despite being a simplification of the real crypt morphology, important biological aspects such as cell heterogeneity and noise are effectively accounted for with this approach.

GRNs are modeled as *Noisy Random Boolean Networks* [27, 28], a simplified model of gene regulation that allows to relate the processes of cell differentiation with the robustness of cells against biological noise and perturbations [29]. This approach focuses on the *emergent behavior* of gene networks in terms of *gene activation patterns* that characterize the cellular activity. Along the lines of [29], each cell type is characterized by particular patterns, whose stability with respect to biological noise is related to its degree of differentiation [30–33]. The approach is general (i.e. it is not related to a specific organism) and is able to reproduce key phenomena of the differentiation processes such as: (*i*) *hierarchical differentiation*, i.e. from toti-/multi-potent stem cells to fully differentiated cells through intermediate stages; (*ii*) *stochastic differentiation*, i.e. a stochastic process rules certain fate decisions and directions; (iii) *deterministic differentiation*, i.e. specific signals or mutations trigger certain differentiation fates; (iv) *induced pluripotency*, i.e. fully differentiated cells can return to a pluripotent stage through the perturbation of some key genes [34].

In our multiscale approach, the GRN dynamics drives cellular growth and the differentiation fate of cells, thus linking the GRN to the crypt morphology.

Following the works by Wong *et al*. [17] and Buske *et al*. [19], in this paper we investigate key dynamical properties of crypts and, in particular, we show that the stochastic differentiation process is itself sufficient to ensure the crypt homeostasis, under certain conditions. Our novel approach permits to relate the genotype-level model of GRN to complex phenotypes and quantitative measures of crucial phenomena occurring in crypts, such as: (*i*) the spontaneous sorting and segregation of cell populations in different compartments, driven by cell adhesion processes; (*ii*) the maintenance of the correct proportion between cell populations with distinct functions in the crypt; (*iii*) the fast renewal process of cells, as resulting from the interplay involving newborn cells and dead ones (either because expulsion in the lumen or apoptosis, which should be modeled per se, cfr. [35]); (*iv*) the coordinate migration of cells from the stem cell-niche toward the intestinal lumen at the top of the crypt; (*v*) the noise-driven progressive differentiation of totipotent stem cells in 8 hierarchical cell types, through transit amplifying stages; (*vi*) the clonal expansion of sub-populations deriving from single progenitors. Furthermore, the model allows to investigate the repercussion of different kinds of gene-level perturbations on the overall dynamical behavior, with a obvious reference to the emergence of cancer.

In this regard, the *dynamical* characterization of genotypic and phenotypic phenomena recently gained greater attention [36, 37]. For example, in [38] cancer development is depicted as a dynamical processes characterized by metastable states (i.e. *attractors* in the terminology of *dynamical systems*) in which stochastic transitions account for cancer heterogeneity and phenotypic equilibria. In general, a dynamical approach provides more information than the static counterpart, given the inherently *evolutionary* nature of cancer. In this respect our model is, to the best of our knowledge, the first attempt to combine a dynamical attractor-based model of GRN with a morphological multicellular model, allowing for innovative analysis perspectives.

Besides, our model is conceived to be flexible and modular, thus both its spatial and gene-level components may be refined to include, for instance, signaling pathways and chemical gradients. We also remark that our modeling approach is general and, in principle, can be applied to any kind of tissue, organ or organism.

The paper is structured as follows. A brief overview of the biology of the crypts is given in the next section. Next, the internal and external components of the model are described, as well as their multiscale link. The results of the analyses on the model are discussed in the subsequent section. Finally, conclusions are drawn.

### A brief overview of the biology of the intestine

Among many, the main functions of the human intestine are (*i*) *food digestion* and (*ii*) *nutrients absorption*, while several other minor processes are linked to the general homeostasis of the system and to the immune system mechanisms. The distinct compartments of the intestine are composed by muscular, stromal and cuboidal epithelial cell. The lining of the small intestine is composed by a single-layer epithelium that covers the villi and the crypts of Lieberkühn, which are the object of our model. Notice that in the large intestine there are no villi, but only crypts (see [1, 39] and references therein).

Four distinct differentiated epithelial cell types are present in the crypt, all descending from multipotent *stem cells*, which give rise to a progeny that undergoes a post-mitotic progressive differentiation process, characterized by the presence of partially differentiated cells in *transit amplifying stages* (see [40] for an exhaustive discussion). In particular, the four epithelial fully differentiated lineages are: *enterocytes*, performing both absorptive and digestive activity via hydrolases secretion, *Goblet cell*, secreting mucus to protect the absorptive cells from digestion, *Paneth cell*, performing defensive tasks by means of antimicrobial peptides and enzymes and *enteroendocrine* cells (a general category with 15 subtypes) entailed in many different tasks and signaling pathways [39, 41, 42]^1^.

Cell type populations are segregated in distinct portions of the crypt: the proliferative cell compartments is in the lower part of the crypt, all other types but Paneth cells reside at its top. Stem cells are sited at the bottom of the crypt in a specific niche, intermingled or just above Paneth cells, according to different hypotheses [44] (see Figure 1). The overall dynamics is a coordinated upward migration of enterocyte, Goblet and enteroendocrine cells from the stem-cell niche [45]. At the end of migration these cells are shed into the intestinal lumen; this loss of cells balances the production from the base of crypt. Paneth cells are the only cells that move downward and reside at the bottom of the crypt (see [2, 40] and references therein). In this complex coordinate movement cell populations maintain the segregation in distinct compartments [1].

**Figure 1.**
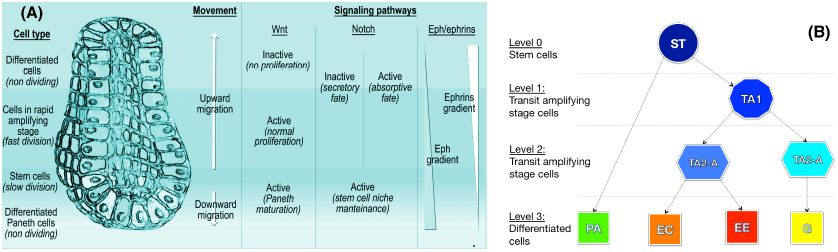
Crypt morphology and differentiation tree. **(A)** A depiction of the crypt morphology, with the direction of cell migration and a schematized representation of the interplay among the key signaling pathways (taken from [13]). All cells but stem and Paneth migrate upward. The three major signaling pathways involved in the crypt activity are the Wnt, the Notch and the Eph/ephrins pathways. In **(B)** the crypt differentiation tree is shown, involving stem (St), transit amplifying stage (TA1, TA2-A, TA2-B), Paneth (Pa), Goblet (Go), enteroendocrine (Ee) and enterocyte (Ec) cells.

The cellular turnover is fast. For instance, in mice the crypt progenitors divide every 12 *÷* 17 hours, so around 200 *÷* 300 cells per day are generated, and they successively undergo up to five rounds of cell division while migrating upwards [39, 46]. Accordingly, migrating cells move from the base to the surface in about 3 *÷* 6 days, while Paneth cells, which live for about 3 *÷* 6 weeks, and stem cells localize at the crypt bottom and escape this flow [2, 47].

The signaling pathways throughout the epithelial cells and between the epithelium and the mesenchyme are fundamental for many phenomena such as spatial patterning, proliferation in transit-amplifying compartments, commitment to specific lineages, differentiation and apoptosis [39]. We briefly describe the three most important signaling pathways involved in these processes.

The *Wnt* pathway is supposed to drive cell proliferation and to rule the differentiation fate. Also, it is responsible of avoiding the immediate differentiation, and activates the expression of the Notch pathway [39]. The activation of this pathway keeps the crypts in a normal proliferative state, whereas its inactivation stops the division/differentiation process. In [44, 48] it is shown that its correct activation is required to determine the Paneth cell fate and lineage.

The *Notch* pathway is involved in the control of the spatial patterning and the cell fate commitment, with the task of ensuring the status of undifferentiated proliferative cells in the progenitors compartment, in a concerted combination with the Wnt pathway [49]. This signaling pathway mediates also *lateral inhibition*, which forces the cells to diversify: some cells express Notch ligands and activate the Notch signaling in the neighbors, while avoiding their own activation. In this way they commit to the finally differentiated fate. In the other cells the Notch ligands are inhibited while the Notch pathway is active within the cell itself; in this way they maintain the possibility of differentiating in any possible way. Multi-potent crypt progenitors are supposed to be maintained only when both Wnt and Notch pathways are active [43].

Finally, the interaction between *Eph* receptors and *ephrin* ligands can trigger a downstream cascade that controls cell-cell adhesion, cell-substrate adhesion, cytoskeletal organization and cell-extracellular matrix binding, influencing the formation and the stability of tight, adherence and gap junctions and integrin functions [50–53].

## A multiscale model of intestinal crypts dynamics

We separately introduce all the model components with respect to the key biological processes we account for. A detailed mathematical definition of the model can be found in the Appendix.

### Crypt morphology as a collective multi-cellular dynamical structure

We adopt a simple geometrical representation of crypts inspired by the theory of cellular automata and statistical physics: the *Cellular Potts Model* (CPM, [24]), often used to account for energy-driven spatial patterns formation [26]. A graphical representation of the CPM model is shown in Figure 2.

**Figure 2.**
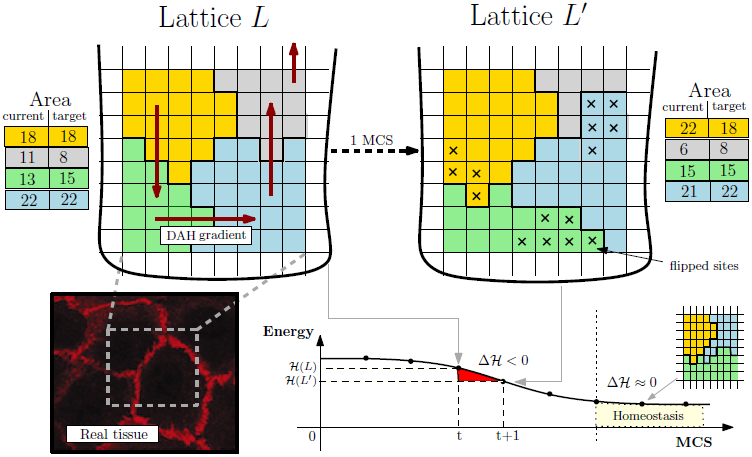
Cellular Potts Model. Lattice-based representation of the crypt tissue as a opened and rolled out lattice *L*, with 4 cells. The energy gradient induced by the DAH via **J** and the current/target area for each cell are represented. An example MCS step is shown resulting in the re-arrangement of *L* in favor of *L*′ (15 flips accepted), whose hamiltonian energy is lower. The final tissue stratification is achieved when Δ𝓗 ≈ 0 where the grey cell is expelled in the lumen. In the left corner an example picture of real tissue is displayed.

We display cells over a rigid 2D grid by assuming a (simplified) perfectly cylindrical crypt, opened and rolled out onto a rectangular *h × w* lattice *L* through periodic boundary conditions. Each cell is delimited by connected domains as in cellular automata so a cell *c*, denoted as *C*(*c*), consists of all lattice sites of *l ∈ L* with value *c*, that is

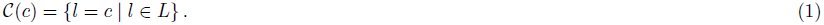

For each disposition of cells a *energy level* is evaluated via a Potts-like Hamiltonian function 𝓗: *L →* ℝ accounting for the energy required for each mutual interaction and other physical quantities (see below). A discrete-time stochastic process of cellular re-arrangement drives the lattice to configurations minimizing the overall hamiltonian energy. The time unit of these steps is the so-called *Monte-Carlo Step* (MCS). The operation key to cellular re-arrangement is that of flipping a lattice site of a cell in favor of another cell, thus modeling cellular movement over the lattice. The changes in the lattice which can happen in a single MCS are sketched as:

1. let *l* be a lattice site selected with uniform probability in *L*, let 𝒩(*l*) be its set of neighbor sites, select a random *l′ ∈* 𝒩(*l*);
2. assign site *l*′ to the cell in *l* with probability

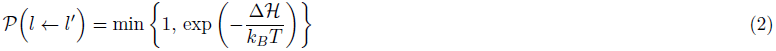

where Δ𝓗 is the gain of energy (i.e., the hamiltonian difference) in accepting the flip;
3. repeat steps 1–2 for *hwk* times, with *k* a positive integer.

In step 1 we set 𝒩(*l*) to the standard Von Neumann neighborhood. Step 2 is the probabilistic re-arrangement of a single lattice site; by iterating *hwk* times a single MCS is simulated and the new lattice configuration displays the cells which moved in that time unit. The Boltzmann distribution is used in equation (2) to drive cells to the configuration with minimum energy; such a distribution depends on the temperature *T* and on the Boltzmann constant *k*_*B*_ (the factor *k*_*B*_*T* gives account of the amplitude of the cell membrane fluctuations at boundaries).

Cell sorting is the phenomenon by which population of cells of distinct type segregate and form distinct compartments or different tissues. According to Steinberg’s *Differential Adhesion Hypothesis* (DAH, [54]), cell sorting may be due to cell motility combined with differences in intercellular adhesiveness and these phenomena in crypts are clearly related to the functioning of the Eph/ephrins signaling pathway (see the Biological background section). In detail, under DAH tissues are considered as vascoelastic liquids whose tissue surface tension can be measured. These tensions correspond to the mutual cellular behavior thought to be responsible for the formation of complex multi-cellular structures. In our model we adopt a *thermodynamical* interpretation of Steinberg’s hypothesis to account for the effects of cell adhesion molecules in a simple way. Along the lines of [17] we assume that a certain amount of energy is required to to keep two cells tied to each other, and we assume that higher energy is required to stick together cells of distinct types. Since the surface tensions can be determined for various tissues, we can use realistic parameter values for these energies [25, 55, 56]. In this way, we implicitly include in our model an abstraction of one of the most important signaling pathways involved in the phenomena relevant to crypt homeostasis.

Therefore, the energy minimized by equation (2) is given by the hamiltonian function

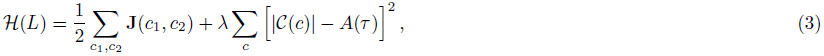

where *c* denotes a generic cell of type *τ* and *c*_1_ and *c*_2_ are different neighbor cells. Function 𝓗 accounts for:

- the amount of energy **J**(*c*_1_, *c*_2_) required to stick tied *c*_1_ and *c*_2_, according to the DAH;
- the tendency of each cell of type *τ* to grow towards some target area *A*(*τ*).

Thus, the target lattice configuration the system is driven to is that where the amount of bond energy is minimal and cells tend to grow up to their target size. Notice that the the total area of a cell is measured as the total number of pixels currently occupied by the cell, i.e., *|C*(*c*)*|*, and the capacity to deform a cell membrane is given by the size constraint *λ* > 0. As far as the DAH is concerned, **J**(*c*_1_, *c*_2_) is the surface energy between the two cells (defined on the basis of the gradients of Eph receptors and ephrin ligands), and is defined according to their cell type (see the Parameters section in the Appendix).

Furthermore, since crypts are not isolated systems, we both consider the expulsion of cells in the intestinal lumen (shedding of fully differentiated cells by mitotic pressure) and the presence of the *Extra Cellular Matrix* (ECM), i.e. the stroma scaffold surrounding crypts. Cell expulsion, which allows the renewal of cells in the crypt, is achieved by the migration of cells towards the top of lattice which, we recall, it is open. The ECM is modeled as a special cell type with un-constrained area (see the Appendix for a detailed definition of function 𝓗 with the ECM cell type).

Finally, cells moving on a lattice eventually complete their cell-cycle. In our case mitosis follows cycle completion and a cell divides into two daughter cells, which are characterized by specific target areas. In particular, stem cells divide asymmetrically, producing a unique daughter (and the stem cell itself), whereas the other proliferative cells divide in two daughters that change type by following the differentiation fate ruled by the GRN dynamics.

### Noise-induced stochastic cellular differentiation via GRNs

We consider the 8 cell types *T* = {St, TA1, TA2-A, TA2-B, Pa, Go, Ec, Ee} shown in Figure 1, and we adopt the hypothesis that *more differentiated cells are more robust against biological noise*, because of more refined control mechanism against perturbations and random fluctuations. Accordingly, the toti-/multi-potent stem cell type is less robust against noise and is thus able to differentiate in any other cell type. In this regard, a wide literature is currently available on: (*i*) the role of noise in gene regulation, e.g., [33, 57–61], (*ii*) the relation between noise and the differentiation processes, e.g., [30, 62–66], (*iii*) the hypothesis according to which the level of noise in undifferentiated cells is relatively higher, e.g., [31, 32, 67].

By using this intuitive idea we link noise-resistance to the *stochastic cellular differentiation* process, at the level of the GRN shared by all the cells in the crypt: once a cell divides, the specific cell type of its progeny depends on a random process, according to the underlying lineage commitment tree. In this paper, we adopt a simplified representation of such a GRN based on the *Random Boolean Networks* (RBNs, [68–70]) approach where genes, and the encoded proteins, are represented in a abstract “on”/“off” fashion. Despite the underlying abstractions, this model has proven fruitful in reproducing several key generic properties of real networks (see, e.g., [71–74]). Intuitively, each gene is associated to a *boolean variable x*_*i*_: *x*_*i*_ = 1, the “on” state, models the activation of the gene (i.e., production of a specific protein or RNA), conversely *x*_*i*_ = 0 models the inactive gene. The interaction among the genes is represented via a *directed graph* where nodes are the binary variables, edges symbolize the regulation paths and each gene affects the neighbor genes via a boolean updating function *f*_*i*_ associated to each node.

The RBN graph represents the possible genetic interactions and is used to “simulate” the evolution of the GRN in a discrete-time, synchronously and deterministically. Let *x*_*i*_(*t*) the state of each gene *x*_*i*_ at time *t*, the new value for the gene at time *t* + 1 is

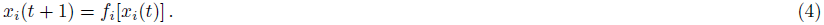

Given that the dynamics is synchronous and deterministic, *gene activation patterns* will eventually emerge from it; technically, these *RBN attractors* are stable *limit cycles* representing sequences of activations/inhibitions of genes, repeating in time [68]. Patterns will be used as a compact representation of the underlying GRN, and their *stability* will be used to model the noise-resistance of each cellular type [27].

This is the so-called *Noisy Random Boolean Networks* (NRBNs, [28, 29]) model of GRN. Together with the DAH-based adhesion energy matrix, this GRN model implicitly includes within the multiscale model the relevant signaling pathways, as their influence is encoded in the various gene regulatory circuits, which, in turn, rule the overall crypt dynamics. We here remark that each cell of the system is characterized by the *same* NRBN, like all the cells of an organism share the same genome (i.e. GRN). The differences in the activity of the distinct cells is due to the particular dynamics of their own gene activation pattern (for instance, distinct cells of the same type own the same NRBN and the same gene activation pattern, but can be in different phases of the pattern).

We sketch here its usage, which is schematized in Figure 3; for a exhaustive mathematical definition of NRBNs we refer to the Appendix. The process is as follows:

**Figure 3.**
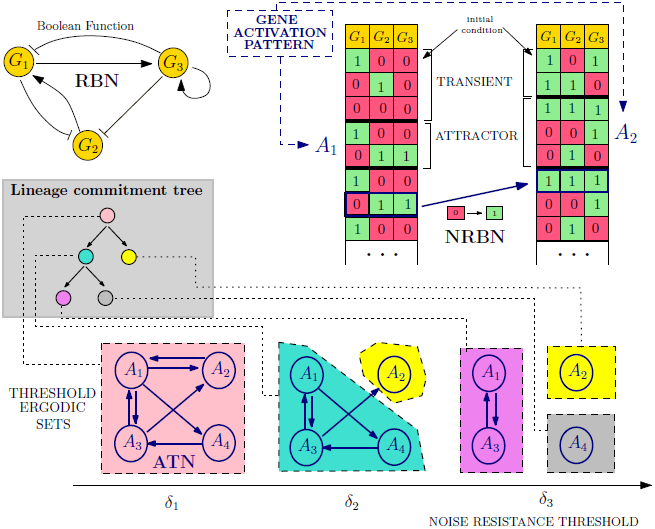
Noise-induced stochastic differentiation. An example NRBN with 3 genes is shown, boolean functions are omitted. Two initial genetic configuration yield two gene activation patterns: attractors *A*_1_ and *A*_2_, whose noise-resistance is evaluated via flipping different nodes in different phases and leading to an Attractors Transition Matrix. The emerging lineage commitment tree consists of 5 cell types (one for each Threshold Ergodic Set for the 3 noise thresholds *δ*_1_, *δ*_2_ and *δ*_3_). The differentiation level corresponds to the noise-resistance, e.g., the toti-/multi-potent stem-alike cell type (pink) roams among all possible gene activation patterns, the grey/yellow cell types are fully differentiated cells.

1. a random RBN is generated with some specific bio-inspired constraints (see below);
2. a set of GRN configurations representing the initial conditions of the RBN is generated by turning “on”/“off” the genes (i.e., assigning 0/1 values to all the variables *x*_*i*_);
3. for each configuration the dynamical trajectory of the GRN is generated via equation (4) (right table, Figure 3);
4. all the stable limit cycles of a GRN define its gene activation patterns (e.g., the attractors *A*_1_ and *A*_2_ in Figure 3);
5. the stability to noise of each gene activation pattern is tested by performing random perturbations on each gene (i.e., temporary *flips*). A stable pattern is robust when the dynamical trajectory that follows a perturbations returns to the pattern itself. Notice that unstable patterns may determine new attractors;
6. by repeatedly performing step (5), the stability of each gene activation pattern is numerically evaluated, determining the noise-induced probability of switching between patterns. The *Attractor Transition Network* (ATN, [29]) accounts for the relative probabilities of switching among patterns (see Figure 3);
7. the *connected components* of an ATN are *noise-driven connected gene activation patterns* used to define the *hierarchical differentiation tree* in Figure 1, more precisely:
  - *toti-/multi-potent stem cells* are the connected component of the ATN involving all the possible genetic patterns, through which the GRN continues to wander due to biological noise and random fluctuations;
  - according to the hypothesis that more differentiated cells are characterized by a higher resistance to noise, we define *threshold-dependent ATNs* by pruning the probabilities below distinct thresholds, hence neglecting the transitions that are unlikely to occur in the lifetime of a cell: higher thresholds correspond to a better resistance against noise. By performing this step recursively, we detect connected components of patterns in the ATN according to increasingly larger thresholds, termed *Threshold Ergodic Sets* (TESs) in the NRBN jargon, which are hierarchically assigned to the sub-types in the tree, according to the strategy defined in [29] (see bottom of Figure 3). Larger thresholds progressively determine smaller and more fragmented TESs, which correspond to more differentiated cell types. The TESs reflect the usual assumptions that less differentiated cells, e.g., stem cells, can roam in the wider portion of the space of plausible genetic configurations for a cell (i.e., *A*_1_, *A*_2_, *A*_3_ and *A*_4_ in Figure 3) and vice versa [61].

When all these steps are complete, the emerging hierarchy between the cell types is matched against the crypt differentiation tree of Figure 1, as sketched in Figure 3. If it matches, the generated NRBN is a network whose emergent cellular types are able to characterize the crypt lineage commitment tree and can be used in the CPM simulation. If it does not match, the NRBN is rejected and the process re-starts.

This strategy requires only a few *a priori* structural assumptions on the underlying GRN, along the usual *ensemble* approach to complex systems [70]. This makes sense since, in this case, it is undoubtedly difficult and hazardous to conjecture a specific human GRN. Instead, we aim at studying the general emergent properties of a class of networks and relating them to the crypt dynamics. In this respect, we generate NRBNs satisfying the structural constraints given by the current biological knowledge of real GRNs and select the *suitable* ones on the basis of their emergent dynamical behavior (see the Results section). Notice that, in line with the fact that the human GRN is unique, we should not expect to find many “suitable” NRBNs.

### A multiscale link between GRNs and the morphology of the crypt

Each cell on the spatial model incorporates a specific GRN, which is characterized by specific gene activation patterns, related to the degree of differentiation. Three major cellular processes are then ruled by the internal NRBN dynamics, thus providing the link between GRNs and the CPM: (*i*) the length of the cell cycle, proportional to the weighted length of the gene activation patterns of each specific cell type, (*ii*) the cell growth rate, assumed to be linear in time, and (*iii*) the differentiation process, as explained in the previous section. We remark that, without accounting explicitly for GRNs, (*i*) and (*iii*) could not be emergent properties but should be assumed.

#### Cell cycle length and time-scales conversion

TESs in the terminology of [29] are analogous to *ergodic* discrete-time Markov Chains, which are known to possess a *unique* computable stationary probability *π* (see the Appendix). We exploit this to evaluate the probability that a cell will be in a certain genetic activation pattern, in the long run. By this, we can infer a measure of the average time needed to reach a stable GRN configuration, thus estimating the cell cycle length.

In formulas, if *π*(*α*) is the stationary probability of a pattern *α*, we define the length ℓ*τ* of the cell cycle for a cell of type *τ* as

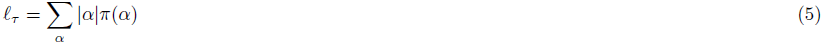

where *|α|* is the number of genetic configurations of the pattern *α* (i.e. number of states of the attractor), which ranges over the set of patterns (i.e. attractors) belonging to the considered TES.

The length of the cell cycle is then an emergent property of the NRBN dynamics, thus a conversion between the involved time-scales is required; this is, to the best of our knowledge, a novel result. We link the internal time-scale (i.e., the NRBN steps) to the external one (i.e., the MCS steps) by considering that (*i*) 10 MCS steps correspond to 1 hour of biological time, according to [17], and that (*ii*) the *average* length of a cell cycle is 12 *÷* 17 hours (we here arbitrarily choose 150 MCS, 15 hours, as a reasonable value to be used in the conversion) (ibidem). Thus, since the natural unit for ℓ_*τ*_ is the NRBN step, we have the following conversion:

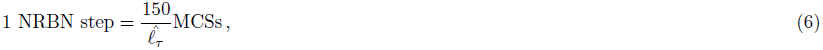

where 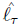 is the *average* cell cycle length of all the cell types of the NRBN. In this way, the relative difference in the lengths of the cell cycles accounts for the difference in the replication pace of the distinct cell types, as a consequence of the emergent dynamics of the GRN. So, for instance, if a cell has only two cell types of length, respectively, 2 and 10, the former type will require 2 *·* 150/ 6 = 50MCSs to complete the cell cycle, whereas the latter will require 10 *·* 150/ 6 = 250MCSs.

#### Cell size dynamics

As we stated above, each cell of type *τ* grows towards a target area *A*(*τ*), and newborn cells have assigned area *A*(*τ*)/2 so they need to double their size before performing mitosis. To spontaneously drive a cell to double its size we make the target area to be time-dependent on the time-scale of the internal GRN, denoted *A*(*τ, t*). As if it was *mechanically isolated*, the time-dependent area grows linearly when the cycle starts at some time *t*_0_, that is

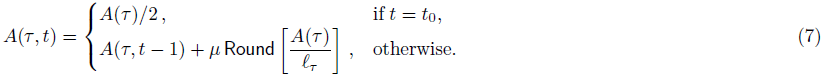

Here we discriminate among proliferative (*µ* = 1) and non-dividing (*µ* = 0) cells; with reference to the tree in Figure 1, non-dividing cells are paneth, goblet, enteroendocrine and enterocyte. Also, Round denotes the nearest-integer function. By introducing this time-dependent area we refine the constraint area term of Equation (3) to be *|C*(*c*)*| − A*(*τ, t*), where *t* is time passed since the beginning of the cell cycle for cell *c*.

#### Cell division and differentiation dynamics

As long as the CPM dynamics goes on, so does the underlying GRN dynamics within each cell, in terms of dynamical evolution of the gene activation patterns. We hypothesize the existence of a certain level of biological noise and random fluctuations, which induces a number of gene mutations: the mutation rate *m* defines the frequency of single flips of genes (as when computing the TESs) and is derived from experimental evidences [75]. In this way, cells that are characterized by TESs with more than one attractor may wander through the distinct gene activation patterns, by means of random mutations.

When a cell concludes in ℓ_*τ*_ NRBN time-steps its cycle and reaches its target size *A*(*τ*) on the CPM, it instantaneously divides and differentiates.

As explained in the previous section, once cells differentiate they increase their noise resistance threshold [29]. The differentiation branch depends on the dynamics of the underlying GRN, as previously discussed and, in particular on the specific gene activation pattern in which the cell is located when the cell divides. Notice that stem cells perform *asymmetric cell division* to preserve their niche, i.e., only one daughter cell differentiates, the other one remains a stem cell [1].

## Results

Simulations of the model were performed by a ad-hoc Java implementation developed by our research group. The search of the NRBN matching the tree in Figure 1 was performed by using GESTODIFFERENT, a CYTOSCAPE [76] plugin to generate and to identify GRNs describing an arbitrary stochastic cell differentiation process [77].

Most of the parameters of the model are set on the basis of experimental data on *mice* and on the general biological knowledge concerning intestinal crypts, whereas the remaining ones are estimated to fit the overall dynamics, with regard to both the spatial and the GRN models. The table with the parameters used in the simulations is reported in the Appendix.

### Properties of the suitable GRNs

As mentioned above, the number of NRBNs with emergent behavior coherent with the crypt lineage commitment tree must be low. Further, no constructive approach is known to determine such networks, and a generative approach is then required.

We here limited our search to NRBNs with certain *structural features* (summarized in the Parameters section in the Appendix) known to be plausible for real GRNs. In particular, we used *scale-free* topologies [78], i.e. NRBNS where the fraction of genes with *k* outgoing connections follows *k*^−*γ*^ for large *k*. Here we used *γ ≈* 2.3 estimated to be a realistic value for many biological networks, including GRNs [79]. The number of genes that we used is 100, which is in line with current belief about the number of genes *relevant* for the correct functioning of crypts, with an average connectivity *|K|* = 3. Finally, concerning boolean functions, we used biologically plausible *canalizing functions* [80, 81].

Our results confirm that finding suitable NRBNs is indeed hard: only 16 out of 4 *×* 10^4^ (i.e. *≈* 0.04%) distinctly generated networks are amenable at use. This confirms that even rather small networks can display a broad range of dynamical behaviors, thus finding the correct emerging lineage commitment tree is hard. This outcome also points to a strong selection process at the base of the emergence and evolution of the current human GRNs.

We tried to *statistically discriminate* among these NRBNs by evaluating some classical measures in network analysis: the number of emerging activation patterns, the average number of genetic configurations they contain, the clustering coefficient of the network, its diameter, the average path length and the average bias of the boolean functions. These measures have been widely used in the characterization of both artificial and real networks such as, e.g., protein-protein interaction networks, GRNs, power grids, co-authorship networks, to name but a few (see [78] and references therein). These statistics are shown in Figure 4. Even if the number of suitable NRBNs is too limited to draw definitive conclusions, the plots hint at the lack of appreciable differences among the suitable and unsuitable networks. Further, this suggests that identifying some GRN parameters to improve this generative approach is indeed hard, as expected by considering that real GRNs are the result of a Darwinian selection process which selected the fittest networks in terms of robustness, evolvability and adaptability to dynamic environmental conditions.

**Figure 4.**
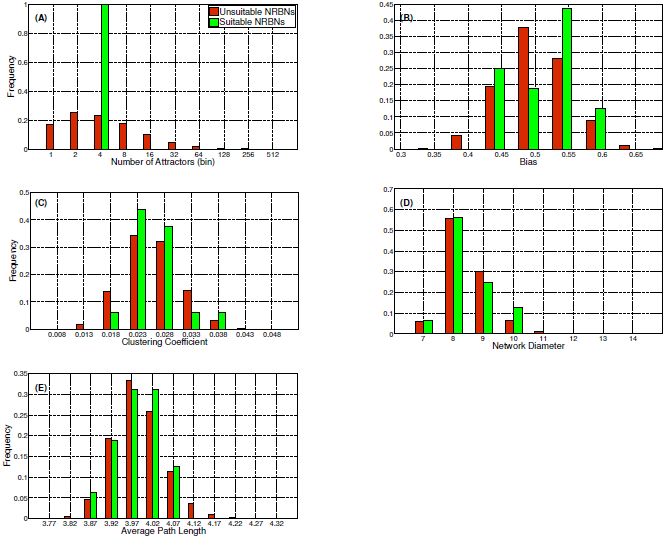
Properties of the generated NRBNs. Statistics regarding: **(A)** the number of emerging activation patterns, i.e. attractors, **(B)** the average bias of the boolean functions, **(C)** the clustering coefficient, **(D)** the diameter and **(E)** the average path length of the generated NRBNs. The distributions are computed on 40.000 simulated NRBNs. The suitable networks are 16. In Tables 1 and 3 we summarize some other key properties of the suitable networks.

**Table 1.** Cell cycle length, in NRBN steps, as computed with equation (5) for the 7 suitable GRNs used in the simulations.

| $\ell_\tau =$ | *type* | *Net*<sub>1</sub> | *Net*<sub>2</sub> | *Net*<sub>3</sub> | *Net*<sub>4</sub> | *Net*<sub>5</sub> | *Net*<sub>6</sub> | *Net*<sub>7</sub> | **Average** |
| --- | --- | --- | --- | --- | --- | --- | --- | --- | --- |
|  | ST | 5.47 | 6.50 | 6.17 | 1 | 20 | 7.23 | 4.35 | 7.24 |
|  | TA1 | 6.60 | 7 | 5.62 | 1 | 20 | 9.8 | 4.81 | 7.73 |
|  | TA2-A | 8 | 7 | 4 | 1 | 20 | 7.50 | 6 | 7.64 |
|  | TA2-B | 6 | 7 | 14 | 1 | 20 | 13 | 2 | 9 |
|  | PA | 4 | 1 | 2 | 1 | 20 | 4 | 2 | 4.86 |
|  | GO | 6 | 7 | 14 | 1 | 20 | 13 | 2 | 9 |
|  | EC | 8 | 7 | 4 | 1 | 20 | 7 | 6 | 7.57 |
|  | EE | 8 | 7 | 4 | 1 | 20 | 8 | 6 | 7.71 |
| | $\hat{\ell}_\tau$ | 6.58 | 6.07 | 5.68 | 1 | 20 | 7.97 | 4.45 | |

**Table 2.** Parameters of the Noisy Random Boolean Networks modeling the Gene Regulatory Network of intestinal crypts, and of the Cellular Potts model of crypt morphology. Parameters with a star symbol are fit.

| Noisy Random Boolean Networks |  |  |  |
| --- | --- | --- | --- |
| Symbol | Value | Description | Source |
| N | 100 | Number of genes (nodes) | *relevant genes for crypt activity (estimated)\** |
| $\vert K\vert $ | 3 | Average connectivity | *input lineage tree\** |
| - | *scale-free* | Network topology | [78] |
| $\gamma$ | 2.3 | Exponent of the power-law of the scale-free model | [79] |
| - | *canalyzing* | Type of boolean functions | [81] |
| Cellular Potts Model |  |  |  |
| Symbol | Value | Description | Source |
| - | $1\mu m = 1 \text{ pixel}$ | Conversion from pixels (side) to $\mu m$ | [6, 19] |
| - | $10 \text{ MCS} = 1 h$ | Conversion from MCS to *hours* | [17] |
| $m$ | 1 flip/MCS | Mutation (single flip) rate | [75] |
| - | $1\text{NRBN}=150/\hat{\ell}_\tau$<br>MCS | NRBN/MCS time conversion | [1] |
| h | $150\mu m$ | Height of the lattice | [6, 19] |
| w | $100\mu m$ | Width of the lattice | [6, 19] |
| k | 4 | Number of lattice spin attempts per lattice site per MCS | *cell turnover\** |
| $\mathcal{N}$ | 7 | Von Neumann neighborhood size | *cell sorting\** |
| $n$ | $\{0, 0.1, 0.25\}$ | Disorder of cellular displacement in the initial configurations | *parameter scan* |
| $\lambda$ | 100 | Area constraint | *cell boundary crumpling\** |
| $k_B T$ | 50 | Temperature and Boltzmann constant | *cell boundary crumpling\** |
| $A(\tau)$ | 50 pixels | Target area for all the cell types | [1] |
| - | 250 MCS | Apoptosis time for Paneth cells | *cell turnover\** |
| **J** | see below | Cell adhesion matrix | [17, 25, 55, 56] |

**Table 3.** Properties of the suitable NRBNs used in the simulations.

|  |  |
| --- | --- |
| $\left( \begin{array}{c\vert cccc} \mathbf{Net_1} & \alpha_1 & \alpha_2 & \alpha_3 & \alpha_4 \\ \hline \vert \alpha\vert & 4 & 6 & 8 & 8 \\ \mathcal{A}(\alpha, \alpha) & 0.985 & 0.956 & 0.812 & 0.825 \\ \pi(\alpha) & 0.456 & 0.351 & 0.102 & 0.09 \end{array} \right)$ | $\left( \begin{array}{c\vert ccccc} \mathbf{Net_2} & \alpha_1 & \alpha_2 & \alpha_3 & \alpha_4 & \alpha_5 \\ \hline \vert \alpha\vert & 7 & 1 & 7 & 7 & 7 \\ \mathcal{A}(\alpha, \alpha) & 0.945 & 0.98 & 0.882 & 0.914 & 0.894 \\ \pi(\alpha) & 0.325 & 0.083 & 0.117 & 0.09 & 0.383 \end{array} \right)$ |
| $\left( \begin{array}{c\vert cccc} \mathbf{Net_3} & \alpha_1 & \alpha_2 & \alpha_3 & \alpha_4 \\ \hline \vert \alpha\vert & 4 & 4 & 2 & 14 \\ \mathcal{A}(\alpha, \alpha) & 0.775 & 0.78 & 0.9 & 0.78 \\ \pi(\alpha) & 0.386 & 0.332 & 0.053 & 0.228 \end{array} \right)$ | $\left( \begin{array}{c\vert cccc} \mathbf{Net_4} & \alpha_1 & \alpha_2 & \alpha_3 & \alpha_4 \\ \hline \vert \alpha\vert & 1 & 1 & 1 & 1 \\ \mathcal{A}(\alpha, \alpha) & 0.8 & 0.74 & 0.92 & 0.9 \\ \pi(\alpha) & 0.371 & 0.342 & 0.057 & 0.228 \end{array} \right)$ |
| $\left( \begin{array}{c\vert cccc} \mathbf{Net_5} & \alpha_1 & \alpha_2 & \alpha_3 & \alpha_4 \\ \hline \vert \alpha\vert & 20 & 20 & 20 & 20 \\ \mathcal{A}(\alpha, \alpha) & 0.886 & 0.877 & 0.896 & 0.891 \\ \pi(\alpha) & 0.262 & 0.266 & 0.251 & 0.221 \end{array} \right)$ | $\left( \begin{array}{c\vert cccc} \mathbf{Net_6} & \alpha_1 & \alpha_2 & \alpha_3 & \alpha_4 \\ \hline \vert \alpha\vert & 8 & 13 & 4 & 7 \\ \mathcal{A}(\alpha, \alpha) & 0.732 & 0.821 & 0.975 & 0.791 \\ \pi(\alpha) & 0.245 & 0.176 & 0.358 & 0.219 \end{array} \right)$ |
| $\left( \begin{array}{c\vert cccc} \mathbf{Net_7} & \alpha_1 & \alpha_2 & \alpha_3 & \alpha_4 \\ \hline \vert \alpha\vert & 6 & 6 & 2 & 2 \\ \mathcal{A}(\alpha, \alpha) & 0.793 & 0.856 & 0.87 & 0.86 \\ \pi(\alpha) & 0.26 & 0.327 & 0.199 & 0.213 \end{array} \right)$ | |

As explained in the previous sections, the emergent properties of the GRN are related to some key features of the cell cycle and differentiation processes at the spatial level. In particular, in Table 1 we show the cell cycle lengths, as computed with equation (5) for the 7 suitable GRNs actually used in the simulations.

It is possible to notice that the length of the cell cycle ranges from 1 to 20 NRBN time steps in different nets and that the variance can be dramatically different among nets, ranging from the case of networks in which all the cell types have the same cell cycle length (i.e. same replication pace), to the case of very different lengths (i.e. very different replication paces). By looking at the average values one can see that most of the cell types have a similar cell cycle length, around 7. Considering that in simulations we set 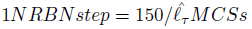, we can the estimate that on average 21 MCS, i.e. around 2 hours, are needed in order to switch among the configurations of a gene activation patterns (i.e. from one state to the following in the attractor). Accordingly, the average cell cycle lasts around 15 hours, which is set to be in accordance with biological knowledge (see the Biological background section). Surprisingly, cell types that are closer in the tree (i.e. Ee and Ec) display almost identical cell cycle length with every network, pointing at an interesting property of such a system.

Distinct other properties of the gene activation patterns of the suitable networks are reported in the Appendix. We here remark that a rather large variability in the robustness to perturbations of the patterns is observed in the different cases, ranging from patterns that are almost imperturbable (99% of the single-flip perturbations end up in the same pattern) to ones that allow switches to other attractors in 30% of the cases after single flip perturbations. This result hints at interesting research perspectives related to the possible advantage for GRN of being *sufficiently* robust to perturbation, while not being too ordered [72]. Furthermore, the analysis of the stationary distributions shows very different scenarios, ranging from the case in which all the patterns are almost equally probable, to that of networks in which some of the patterns are very unlikely (e.g. less than 5%). Also in this case, it would be interesting to match these results against experimental evidences, to investigate the role of the temporal permanence within the same pattern and of the transitions among them.

### Cell sorting and overall homeostasis

The major goal of this work is to determine under which conditions the correct functioning of intestinal crypts is ensured and maintained, with particular reference to *cell sorting, coordinate migration* and *general homeostasis*.

To this end, we analyzed the crypt dynamics via CPM simulation, by using the suitable NRBNs. Please refer to the Parameters section in the Appendix for the parameters of the CPM used in the simulations. To account for the role of the initial displacement of cells within the crypt we tested 4 distinct configurations on a 100x150 pixels lattice, according to the initial level of “order” (we set 1 pixel side to 1*µm*, to agree with experimental evidences [6, 19], see the Appendix). A *disorder* parameter, *n* discriminates the first three configurations: *n* = 0 denotes a configuration in which the cells are perfectly sorted, *n* = 0.1 (resp. *n* = 0.25) a configuration in which 10% (resp. 25%) of the cells are randomly positioned on the lattice. The fourth initial condition is composed only of stem cells, positioned at the bottom of the crypt, while the remaining lattice is empty. The latter configuration aims at investigating *in-silico* the dynamics of isolated stem cell progeny populations, as classically done via *in-vitro* experiments [82].

In all the initial conditions cells are assigned a square shape^2^: in the first three cases 560 cells are displayed with the following cellular proportions: 60 stem cells, 60 Paneth, 240 TA-1, 120 TA-2 and 80 differentiated cells. In the fourth case 120 stem cells are considered. The initial conditions are shown in Figures 5 and 6, together with some sampled crypts after 2000 MCS (200 *h*) with 50 final *annealing steps*^3^. For each of the 7 suitable GRNs we performed 10 independent CPM simulation runs, in order to have a relevant statistics. We remark that the values of **J** are based on experimental results showing that a high activation level of the Eph receptor reduces cell adhesion and vice versa [55, 56] (Parameters section in the Appendix). Only the relative magnitudes of cell adhesion energies are needed to our modeling approach.

**Figure 5.**
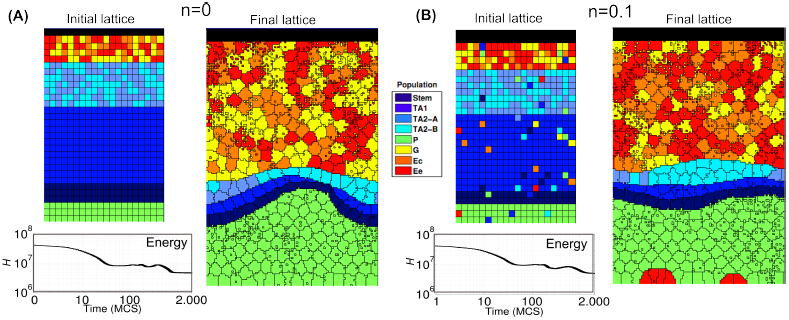
Crypt homeostasis - 1. Initial lattice configurations for *n* = 0 (**A**) and *n* = 0.1 (**B**) and corresponding lattice after simulating 200 hours of crypt evolution, for a single simulation. The overall system energy is the average of 70 independent simulations. Crypt layout was drawn by using the visualization capabilities of CompuCell3D [92].

**Figure 6.**
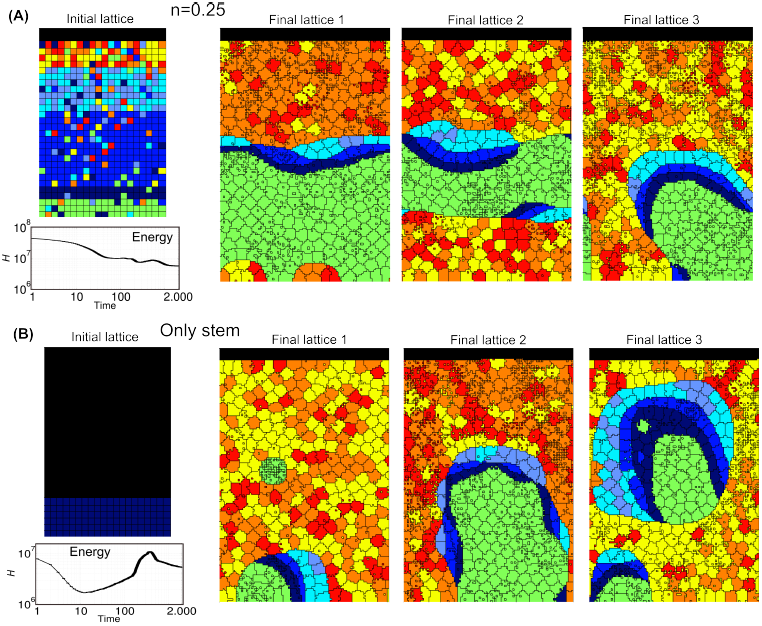
Crypt homeostasis - 2. Initial lattice configurations for *n* = 0.25 (**A**) and the case of only stem cells (**B**), and corresponding lattice after simulating 200 hours of crypt evolution, for a single simulation. The overall system energy is the average of 70 independent simulations.

By these figures it becomes clear that the final crypt ordering is dependent of the initial ordering. In particular, for very low-noise configurations the correct crypt behavior always emerges. Differently, in the case for *n* = 0.25 deeply different scenarios are displayed at each simulation. In some cases, the correct cell stratification is achieved, while in others some distinct geometrical shapes, e.g., encapsulations and invaginations, are observed, and the overall homeostasis is not achieved. In the fourth initial configuration (i.e. only stems), it seems unlikely that the crypt may reach a correct stratification. In the next sections we analyze these scenarios in detail by evaluating specific statistics.

Notice that the overall system energy (i.e. the Hamiltonian 𝓗), whose variation time is shown in the figure, asymptotically reaches a minimum value which ensures an optimal (dynamical) configuration of the cells on the lattice. In the specific case of stem cells (Figure 6), one can observe a peak in the Hamiltionian after around 1000 MCS. This phenomenon is due to the expected progressive appearance of large populations of distinct differentiated types, as opposite to the relatively more favored initial configuration, in which only cells of a unique type (i.e. stem) are present in the system.

One of the most important results of these (and the following) analyses is to show that in our model the stochastic differentiation at the GRN level is itself sufficient to ensure the *normal* activity of the crypt, in terms of overall spatial dynamics. This results is even more surprising by considering that, as shown in the previous section, the lengths of the cell cycles are indeed different in the distinct suitable networks used in the simulations. Hence, it is reasonable to hypothesize the existence of a relatively broad region of the gene activation space in which the correct functioning of the crypt is maintained, despite the differences in the replication pace of different cell types, as long as a suitable differentiation tree is maintained to ensure the correct cell turnover. Besides, with this approach no explicit signaling pathways are considered, which instead result from the interplay between the GRN and the CPM features. Interesting research perspectives derive form this outcome, with particular regard to the configuration of the activation patterns related to the emergence of *aberrant* structures.

### Cell population dynamics

The variation in time of the *number of cells in each population* is shown in Figures 7 and 8 for the four distinct initial configurations. Despite some differences, in all the cases an asymptotic stable proportion is reached, after a transient in which the crypts tend to adjust. In particular, a proportion between the cell types is maintained in all the cases, predicting quantities that are in agreement with what is supposed to be the general proportion of cell populations in real crypts, i.e. around 300 [1, 17, 19, 83]. More in detail, the *average* final configuration involves cell population in these proportions: Stem cells 2.5%, TA1 2.5%, TA2-A 2%, TA2-B 2%, Paneth 27%, Goblet 22%, Enterocite 22% and Enteroendocrine 20%. Surprisingly, this pseudo-equilibrium is reached regardless of the different initial conditions, suggesting that the GRN-driven crypt dynamics is able to ensure a “correct” cellular proportion. The only clear difference predicted by the initial conditions is that, in the case of a crypt with only stem cells, the system appears to have a longer transient.

**Figure 7.**
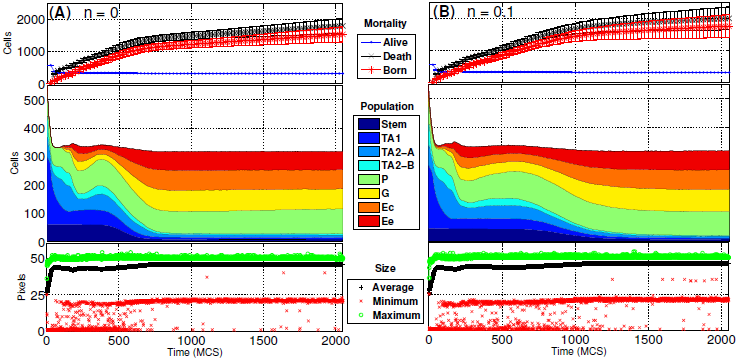
Dynamics of the cellular populations - 1. Number of cells for each cellular population, number of newborn, dead and alive cells and maximum, minimum and average cell size, in time. Notice the prediction of 300 cells, regardless of the two initial conditions *n* = 0 (**A**) and *n* = 0.1 (**B**). The length of the transient is similar, in both cases.

**Figure 8.**
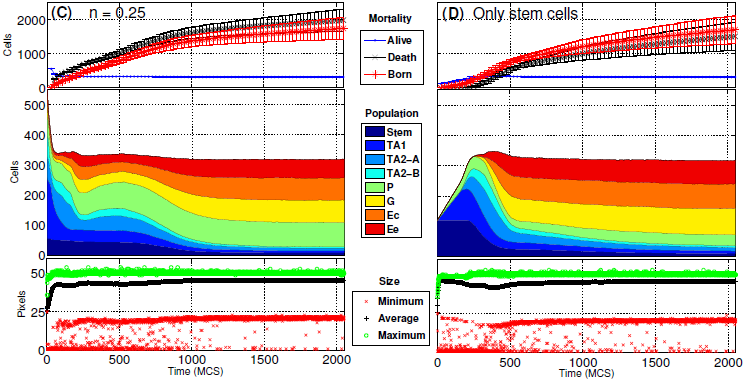
Dynamics of the cellular populations - 2. Number of cells for each cellular population, number of newborn, dead and alive cells and maximum, minimum and average cell size, in time. Notice the prediction of 300 cells, regardless of the two initial conditions *n* = 0.5 (**C**) and the case with only stem cells (**D**), with longer transient.

In the same figures we also show the *number of newborn* and *dead cells* (either due to apoptosis or to the expulsion in the intestinal lumen). Even these two quantities tend to a dynamical equilibrium for all the distinct initial conditions, hinting at an intrinsic capability of the system to ensure a *correct dynamical turnover* or, in other words, the *renewal of the tissue*. The quantities shown in the figures agree with the phenomena supposed to characterize real crypts [17].

Finally, the maximum, minimum and average *size of each cell* are shown. We remark that the initial cells are newborn, so their size is half of their target area when mature, i.e. just before undergoing mitosis. This is the reason why all the initial values of this statistics, particularly in the average case, are much lower than the asymptotic ones which are, in any case, stable. One can see that the average cell size is very close to the maximum, suggesting that the crypt mostly contains “adult” cells. Also, being the variance relatively small, this suggests that cells have similar sizes, on the average. Finally, the maximum size has an upper bound proportional to the pre-mitotic size, which is only rarely exceeded due to random fluctuations.

### Coordinate migration

From experimental results it is known that cells at the bottom of the crypt move slower toward the top than cells positioned in the upper portion [17, 84]. By looking at Figure 9 we can notice that there is a correlation between the distance from the bottom of the crypt and the average *vertical velocity* of cells^4^. We do not show the minimum vertical velocity which is 0, for some cells and we also remark that the average values do not take into account the fact that some cell populations (e.g. stem cells) move much less than others (e.g. differentiated cell), as required. Recalling that 1 pixel side, *p* = 1*µm*, the average vertical velocity of cells ranges from 0 *p/MCS* at the bottom of the crypt to 0.25 *p/MCS* at the top, that is at most 2.5 *p/hour*. Hence, we can estimate the average time needed for a random descendent of a stem cell (and originating in the stem cell niche), to complete the (progressively faster) migration toward the lumen. It turns out that *around* 650 MCS, i.e. 65 hours, around 3 days, are needed and this result is in perfect agreement with experimental data [1, 85]. We can also notice that the maximum observed vertical velocity ranges from around 0 at the bottom of the crypt to around 8 *pixel/MCS* at its top, with regard to all the configurations. This outcome indicates that some cells can move dramatically faster than other in the overall spacial displacement, due to local energy configurations.

**Figure 9.**
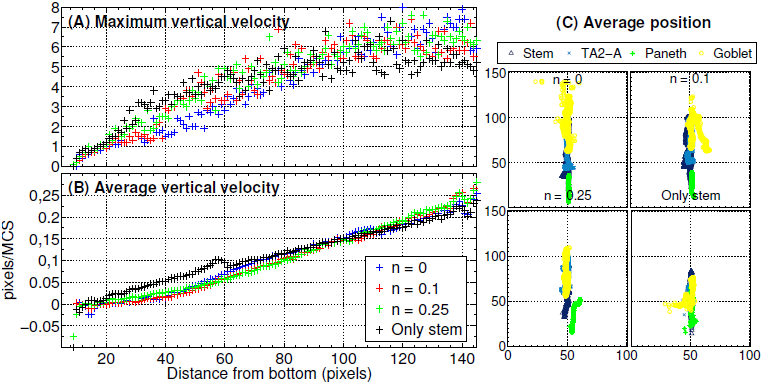
Cell migration. Relation between the distance from the bottom of the crypt (in pixels, i.e. *μm*) and the maximum (**A**) and average (**B**) vertical velocity of the center of mass of all cells. This is averaged for all simulations, at all the time steps. In (**C**) we show the average center of mass for stem, Goblet, Paneth, TA2-A, averaged on all simulations and sampled every 20 MCS.

In order to highlight the relative positioning of cell populations during a simulation, in Figure 9 one can see the movement of the average center of mass of the cells belonging to four distinct types, i.e. stem, Paneth, Goblet and TA2-B, during the whole simulation. A general correct positioning of the populations is maintained with all the distinct initial configuration, yet as long as the level of disorder increases the displacement becomes less precise, as for the case of only stem cells. In any case, a coordinate migration involving the whole crypt is proven to be an emergent property of the GRN-driven dynamics.

This and the subsequent analyses prove that cells translocate in a coordinate fashion towards the top of the crypt, as observed *in-vivo* [85].

### Quantitative measures of spatial ordering

Experimental evidences suggest that epithelial cells migrate in coordination as sheets in culture [86]. Along the lines of [17] we determine whether our cells move coordinately by using the following *spatial correlation index* [86]:

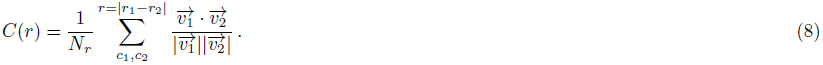

Here *r* is the distance between the center of masses of two generic cells *c*_1_ and *c*_2_, 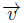 is their cell velocity and *Nr* is the overall number of cell pairs with distance equal to *r*. If a inverse reciprocity relation holds between *C*(*r*) and *r* this implies that closer cells display a more coordinate movement than distant ones. This is what we actually observe in Figure 10: the movement of the cells is highly correlated, unless for very distant cells, which also show large fluctuations. For the stem cells case we observe a slight decrease in the average correlation. This outcome closely resembles the one shown in [17], confirming that the coordinated cellular movement is maintained when also a GRN is used to drive the stochastic differentiation dynamics.

In order to automatize the evaluation of the general *spatial order* of crypts, we propose to use the *Moran Index* (MI, [87]) and the *Pearson’s correlation coefficient* (PC). These measures will allow to understand if the cell populations form groups, and if the the correct stratification is achieved. We recall their definition here, extended to matrices^5^; the PC *ρ* for two matrices **x** and **y** is a function of their (co)variances

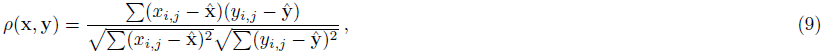

where *x*_*i,j*_ is a component of **x**, and 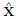 is its average. The PC ranges from *−* 1 (inversely correlated) to 1 (correlated) and at 0 there is no correlation between **x** and **y**.

The PC is also used in the MI, which is used to determine if lattice positions are correlated, that is if cells are likely to form strains of the same type. To define the MI we associate, to each cellular type *τ*, a unique integer value (so 8 values in total), and we evaluate, for each position, the average of all its neighbor cellular types. In formulas, for a position *l ∈ L* we evaluate

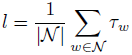

where 𝒩 is the set of neighbors of *l* in *L* (we used the 1^*st*^ order Von Neumann neighborhood), and *τ*_*w*_ is the integer associated to the cell type in *w*. This formula yields a new lattice *L*_𝒩_ to compute the MI as

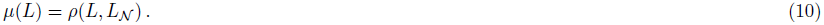

Notice that *−* 1 *≤ µ ≤* 1 with the usual meaning, and that the MI is equivalent for two symmetrical lattices. Thus, despite being a good measure for aggregation, the MI itself does not distinguish if a crypt is stratified with the correct bottom-up ordering, or, for instance, if it reversed. We can anyway use a *template lattice T*, i.e. a lattice were the cellular stratification is made explicit, to asses, the PC between a lattice and the template, that is

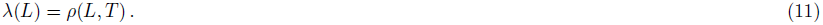

By combining these measures we can state that a crypt is well stratified if it has high *µ*(*L*) (i.e. it has high MI thus it is stratified), and it has high *λ*(*L*) (i.e. it is highly correlated to the template, thus it has the cellular populations correctly stratified).

All these spatial measures are plot in Figure 10. Initially, the MI (averaged over all the simulations) is clearly dependent on the lattice initial condition. After a transient where the stratification level decreases, the MI asymptotically approaches a high value (still proportionally to the initial level of noise), in all but the only-stem-cells case, where the MI gets highly dispersed. This suggests that the stratification is generally maintained, with distinct GRNs and initial conditions, in all cases but when only stem cells are present, a clearly particular scenario.

We compared these statistics with the PC for the corresponding simulations and the template lattice shown in Figure 10. The template lattice considers the proportion among cell populations that is derived from the average final configuration of the correctly stratified lattices (see above). The variation of the PC in time seems to dependent on the initial condition, thus giving further information besides the “general” degree of order depicted by the MI. The PC suggests an inverse proportionality between the initial noise and its asymptotic value, thus hinting at the importance of the initial crypt morphology for its development in the preliminary stages. As for the MI, the lowest PC is for the lattice with only stem cells since we do not impose any constraint on the spatial development of the crypt besides upper/lower bounds. Movement direction and expansion of the cell population emerges from the dynamics induced by the underlying GRN.

**Figure 10.**
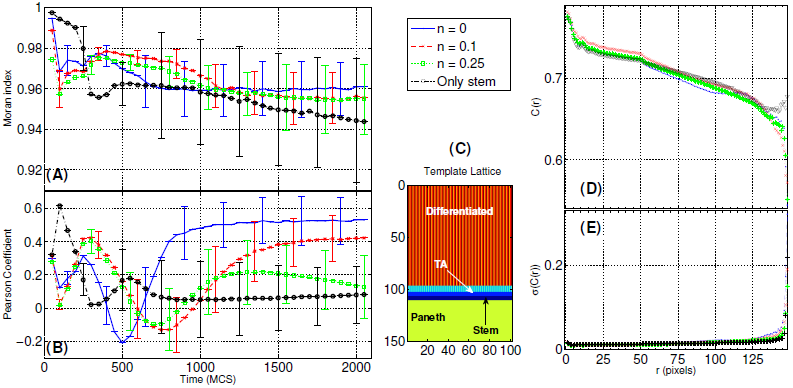
Spatial statistics for crypt stratification and coordinate migration. Time-variation of the Moran Index, MI (eq. 10) (**A**) and of the Pearson Coefficient, PC (eq. 11) (**B**). In (D) the Spatial Correlation *C*(*r*) (eq. 8) with the relative standard deviation (**E**) is displayed, for the four initial conditions. A crypt can be considered well stratified if its MI is high (it is indeed stratified), and its PC is high (it has the cellular populations in the correct order), according to a template (**C**).

### Clonal expansion

Our model implementation permits to track the descendant of each stem cell in the crypt, thus allowing to investigate the process of clonal expansion within intestinal crypts. This will help to determine, in future works, whether any relevant difference is detectable with respect to the case of cancer evolution. It is, in fact, known that tumors develop through a series of clonal expansions, in which the most favorable clonal population survives and begins to dominate, in a Darwinian selection scenario [88].

In Figure 11 we show the distribution of the number of descendants of each *proliferative* cell in the system, for each simulation we performed. Regardless of the initial configuration in most cases 50 *−* 70% of all proliferative cells have a small number of descendants (below 5), whereas only in certain cases a few cells actually display a relatively high number of descendants (almost 80). For instance, in simulation 28 the cell with the largest progeny has 88 descendant, but it actually fails in colonizing the whole crypt or even a significant proportion of it. In fact, at the end of the simulation only around 15 cells out of the 88 descendants are alive thus composing that specific clonal population; these are only the 5% of the overall population (almost 300 cells). This outcome points once more to the homeostasis of the system, in which a *correct* proportion of the cell populations is maintained along the course of the simulation. We expect that serious mutations of the underlying GRN may lead to the appearance of fast replicating clones, which may eventually colonize the whole crypt or a relevant part of it, inducing the emergence of aberrant structures. We finally remark that a larger number of descendants (on average) is observed in correspondence with systems characterized by a higher level of disorder in the initial condition, and this hints at the important role of a correct stratification in maintaining the *right* pace of division in the crypt.

**Figure 11.**
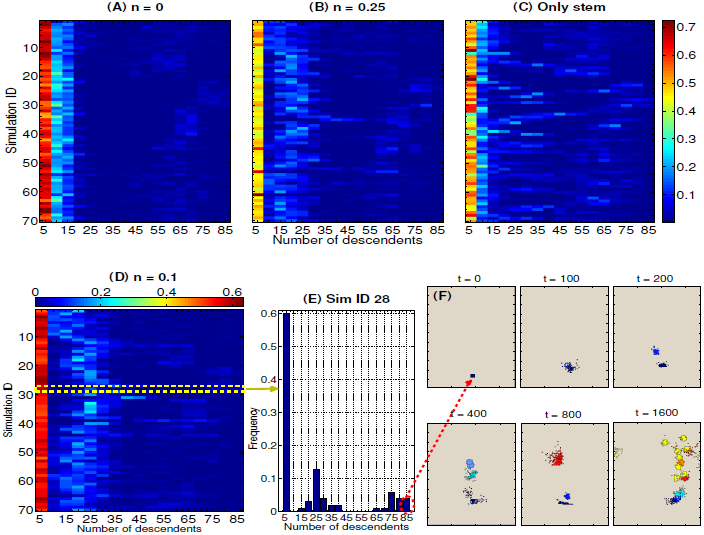
Clonal expansion. Discrete probability distributions (as *heatmaps*) of the total number of descendants of any cell during each simulation with (**A**) *n* = 0, (**B**) *n* = 0.25, (**C**) only stems and (**D**) *n* = 0.1. (**E**) is the histogram for simulation n.28 with *n* = 0.1. In (**F**) we show the alive progeny of that stem cells at the instants *t* = {0, 100, 200, 400, 800, 1600}, colored according to the cell types.

## Conclusions and further development

In this paper we introduced a novel multiscale model of intestinal crypt dynamics, by combining a well known in-lattice model from statistical physics to a boolean GRN model from complex systems theory. This model relies on a few assumptions only, thus reducing the number of its parameters, and the multiscale link between the crypt morphology and its genotype results from the emergent properties of the underlying GRN.

The model allows to efficiently investigate many dynamical properties of crypts such as, e.g., cell sorting, coordinate migration, stem cell niche maintenance and clonal expansion. On the overall, the model suggests that the fundamental process of stochastic differentiation may be sufficient to drive the overall crypt to homeostasis, under certain crypt configurations. Our approach allows also to make precise quantitative inferences that, when possible, were matched to the current biological knowledge.

The model itself was conceived to be flexible and modular, thus all of its components will be possibly refined in future works, along the lines of other approaches (see the references provided in the introduction). In this first paper we focused on studying the development of healthy crypts, and we tried to assess the model conditions under which the activity of a normal crypt emerges and is maintained. These results will be used as a base for future research directions, all of them pointing to multiscale studies concerning the emergence of colorectal cancer, which is supposed to originate in crypts, most likely in the stem cell niche [2].

To this end, the choice of the internal GRN model allows for many possible improvements and research perspectives. For instance, along the lines of the usual NRBN approach, the effect of genetic perturbations of various types (e.g. *gene mutations*) will be assessed with respect to the emergence and development of cancer. Possible communication mechanisms among the GRNs of neighbor cells may be introduced in the model as in, e.g., [89], as well as more accurate descriptions of gene activation and dynamics as in, e.g., [90]. Also, the role of the extrinsic noise in the system, e.g. random thermodynamic and kinetic fluctuations, might be quantitatively assessed as discussed, for instance, in [91].

Furthermore, the networks that we found suitable to describe the lineage commitment tree for crypts will be matched against the currently known portions of the human GRN by employing, for instance, *graph isomorphism* techniques. Also, current knowledge will be used to set up constraints on networks generation, possibly allowing to infer new portions of the human GRN related to the genes involved in the activity of the crypts. To address this ambitious goal, the relevant genes and their interactions involved in the evolution of colorectal cancer could be explicitly considered in the generation of the GRNs to be used in our model.

## Acknowledgments

We thank A. D’Onofrio for suggesting the use of the Moran Index and the Pearson Coefficient as spatial statistics of crypt morphological development. We also thank Silvia Crippa for contributing in developing the software tool for the simulations. This work was partially supported by the ASTIL program, project “RetroNet”, grant n. 12-4-5148000-40; U.A 053 and NEDD Project [ID14546A Rif SAL-7] Fondo Accordi Istituzionali 2009.

## Appendix

### Parameters

In Table 2 we report the parameters used to generate the NRBNs and to set up the CPM simulations. The results of the simulations are discussed in the Results section.

### Mathematical specification of the model

We here present the full mathematical specification of our multiscale model. With respect to the main text some notation might differ, when appropriate.

#### The lattice-based model of cellular tissue

Within a *n × m* 2D *lattice of positions L ∈* {1,…, *k*}^*n×m*^ a population of *k* cells of any type is placed. We denote with *l*_*i,j*_ the single-site variable (spin in CPM terminology) associated with position (*i, j*), we write *l*_*i,j*_ = (𝒸, τ) if the position is occupied by a cell 𝒸 ∈ 𝒞 of type τ ∈ 𝒯, i.e. *T* = {St, TA1, TA2-A, TA2-B, Pa, Go, Ec, Ee} ∪ {Ecm} being the finite set of cell types we consider plus a special type for the ECM. A cell 𝒯(𝒸, τ) with *current area a* (𝒸, τ) is defined by

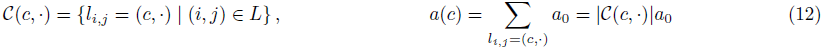

where *a*_0_ is a basic area (in unit of pixels *p*) assigned to a single lattice position (a pixel).

The mutual interaction energy is defined via the Hamiltonian 𝓗: 𝕀^*n×m*^ → ℝ

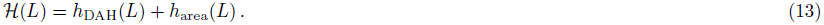

Let 𝒩(*i, j*) denote the Von Neumann neighborhood of (*i, j*), and let *τ*_*i,j*_ denote the type of cell in *l*_*i,j*_, the contribution to the energy according to the DAH is

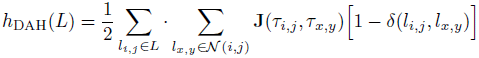

where *δ* is the Kronecker delta function (which ensures that only the surface sites between different cells contribute to the adhesion energy), and **J** is the surface energy discussed in the main text. Let *τ*_*c*_ denote the type of cell *c*, the contribution to the energy of the area constraint is

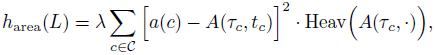

where the ECM has unconstrained area, i.e. *A*(Ecm, *·*) is constant and negative, and its constraint is suppressed by the Heaviside function Heav(*·*). Notice that here we made explicit the time-dependency of the area constraint by using *t*_*c*_, the relative time since the beginning of the cell cycle for *c*.

Given a lattice *L*, we define the flip of a position (*x, y*) to a cell *c* to be the new lattice *L* [*c ←* (*x, y*)] where

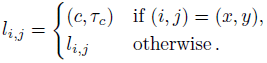

The CPM simulation is as follows: uniformly pick a position (*i, j*) of *L* and pick a neighbor position occupied by a cell *c* (uniformly distributed on the set 𝒩(*i, j*)), i.e. the candidate flip. The CPM probabilistically accepts or rejects the flip, i.e. keeps *L* or update it to *L* [*c ←* (*i, j*)], according to the energy of both lattices and the temperature-dependent probability distribution discussed in the main text, when *k*_*B*_*T* > 0. Instead, when *k*_*B*_*T* = 0 the distribution is

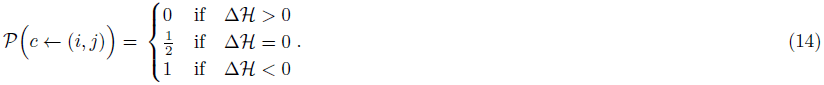

A computation starting at time *t*_*s*_ and ending at time *t*_*e*_ performs *t*_*e*_ − *t*_*s*_ Monte Carlo steps each one attempting *nmk* random flips, as discussed in the main text. Once all the attempts of flips are finished, the new lattice is determined as a result of all the accepted flips. A stochastic process {*L*(*t*) | *t* = 0, 1,…} whose states are the lattice configurations, underlies the CPM. Technically, such a process is a Discrete Time Markov Chain.

#### The boolean model of Gene Regulatory Network

Let **x**(*t*) be the RBN binary vector-state at time *t*, its *i*-th component in **x**(*t* + 1) is

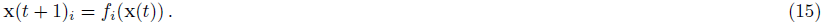

An execution of an RBN is a series of steps from an initial state **x**(*o*)

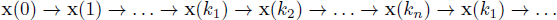

which, since the state-space is finite (i.e. for *w* genes there exist at most 2^*w*^ vectors in {0, 1}^*w*^) and the dynamics is fully deterministic, will enter a limit cycle **x**(*k*_1_) *→… →***x**(*k*_*n*_) from any initial condition. Such a cycle is termed *RBN attractor*, the sub-sequence from **x**(0) to **x**(*k*_1_) (excluded) is its *transient*; the set of initial conditions ending up to the same attractor is its *basin of attraction*.

Our approach first selects, either exhaustively or randomly, a set *I* = {**x** ∈ {0, 1}^*w*^} of initial conditions, and determines numerically its associated set of attractors *A*_*I*_. Iteratively, in each attractor in *A*_*I*_ random flips are performed, i.e. some **x**_*i*_ is complemented in {0, 1} yielding a new state **x**′. The attractor *α* reachable from **x**′ is then computed and included in *A*_*I*_, if *α* ∉ *A*_*I*_. By an arbitrary iteration of this process the ATN 𝒜 of the subsumed NRBN is drawn as follows: for each perturbation yielding a jump among attractors *α* and *β* the entry *a*_*α,β*_ ∈ 𝒜 is incremented, the matrix is finally normalized. As running example, consider the following ATN

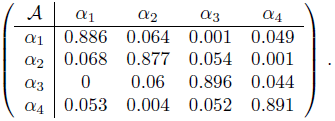

The differentiation tree *T* induced by 𝒜 is generated according to the strategy discussed in [29]. With no aim of exhaustivity we briefly recall it here. Firstly, 𝒜 is interpreted as the *adjacency matrix* of the graph of the attractors *G*_𝒜_. Secondly, the set of *strongly connected component* in *G*_𝒜_ is evaluated by standard algorithmic techniques; this is the TES at threshold 0, *T*_0_, which is supposed to contain all the nodes of *G*_𝒜_. If *T*_0_ does not contain all the required nodes, the RBN is rejected and the process restarts with a new network. Otherwise, *T*_0_ is assigned to the root of the differentiation tree. In the above ATN, *T*_0_ = {(*α*_1_, *α*_2_, *α*_3_, *α*_4_)}.

This process now iterates proportionally to the size of the tree we want to generate, which has a maximum theoretical depth proportional to the number of nodes when *T* reduces to a path. The idea is that a threshold 0 *< δ <* 1 is picked, 𝒜 is pruned of any *a*_*α,β*_ *< δ*, and the new number of TESs at threshold *δ, T*_*δ*_, is evaluated. The set of direct descendants of the root of *T* is defined by *T*_*δ*_ via an injective map, i.e. each TES in *T*_*δ*_ defines a unique descendant. In the example above, when *δ* = 0.053 the pruned ATN is (in bold the pruned entries)

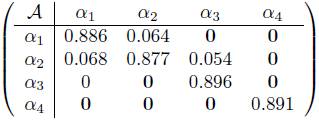

which induces *T*_0_ = {(*α*_1_, *α*_2_), (*α*_3_), (*α*_4_)}. The process stops when no more TESs emerge, and the last descendants determined become the leafs of *T*. For the example above further candidates thresholds are 0.06 and 0.068.

#### The multiscale link

Each 𝒜, re-normalized once a threshold is fixed, is actually a Discrete-Time Markov Chain (DTMC) and the sub-matrix 𝒜_*θ*_ associated to the emerging TESs *θ* is, by definition, a ergodic DTMC (i.e. it is possible to go from each state, in any number of steps, to every other state). In DTMCs terminology the transition probability matrix 𝒜_*θ*_ is *irreducible* and thus the *stationary probability distribution* of the chain

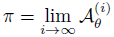

is unique and can be evaluated as the fixed point of the recursive transformation

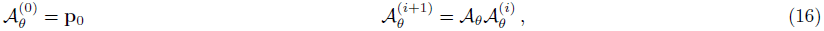

where **p**_0_ is an arbitrary initial distribution over the states of the chain. By fixing an arbitrary small *∈*, this fixed point can be evaluated yielding an approximation *π*\* ≈ π, which can be used as explained in the main text.

### Networks analysis

In our model GRNs drive the spatial dynamics, thus we analyzed different properties of the networks matching the crypt lineage tree. In table 3, one can find some statistics for the 7 suitable network used in the simulations: (*i*) the length of any single attractor of the network (i.e. the number of states of all the gene activation patterns) *|α|*; (*ii*) the robustness of any attractor, in terms of probability of remaining in the same attractor after one single flip perturbation, which is the first entry of ATN matrix, *A*(*α, α*); (*iii*) the stationary distribution of the ATN for each attractor *π*(*α*), which accounts for the average portion of time (in percentage) spent in each attractor and which is used to weigh the length of attractors when computing the cell cycle length. Please refer to the Results section for the comments on these measures.

## Footnotes

1 Other minor cell types, such as M-cells and Brush cells have been also detected [43].

2 Clearly, the initial square shape of the cells is a strong simplification, which however does not affect our analysis, because the energy minimization-driven dynamics leads the cells to more physically plausible shapes in a few MCSs.

3 It is known that, by performing simulations at nonzero temperature, cells are not required to be connected and cell boundaries can crumple, especially when the temperature is comparable to the boundary energy. Glazier and Graner suggest to use a certain number of zero-temperature annealing steps to remove these defects, even if this procedure evolves the spatial pattern as well [25]. Nonetheless, we here remark that this kind of lattice artifacts are not relevant to our analysis, which is based on the statistical analysis of quantitive measures at a coarser grain.

4 Only the *vertical component* of the velocity is here shown, the positive values being associated to the direction toward the top of the crypt.

5 Originally, these measures are defined for vectors however we will use them for our lattices, which are matrices. This extension is straightforward.

